# The Resistome: updating a standardized resource for analyzing resistance phenotypes

**DOI:** 10.1101/418814

**Authors:** J.D. Winkler

**Keywords:** synthetic biology, standardization, adaptive evolution, genomics

## Abstract

Advances in genome engineering have enabled routine engineering and interrogation of microbial resistance on a scale previously impossible, but developing an integrated understanding of resistance from these data remains challenging. As part of our continued efforts to address this challenge, we present a significant update of our previously released Resistome database of standardized genotype-resistance phenotype relationships, along with a new web interface to enable facile searches of genomic, transcriptomic, and phenotypic data within the database. Revisiting our previous analysis of resistance, we again find distinct mutational biases associated with random selection versus genome-scale libraries, along with pervasive pleiotropy among resistant mutants. Attempts to predict mutant phenotypes using machine learning identified the lack of comprehensive phenotype screening and small size of the Resistome corpus as challenges for effective model training. Overall, the Resistome represents a unique platform for understanding the interconnections between both current and future resistant mutants, and is available for use at https://resistome-web-interface.herokuapp.com.

## Introduction

Microbial stress resistance has significant implications for human health and industrial biotechnology. For example, antibiotic treatments may fail as resistance-encoding elements are acquired via horizontal gene transfer or evolution (*1*) and biocatalysts should endure extremes of pH, temperature, osmotic strength, and product toxicity in industrial fermentation (*2, 3*). While decades of research have provided the tools to examine the molecular mechanisms underlying genotype-phenotype relationships, our ability to generate and characterize resistance mutants via selection (*4*–*8*) and screening genome-scale libraries (*9*–*12*) has outstripped improvements in our ability to leverage these voluminous datasets for forward engineering or prediction of resistance. Leveraging the accumulating biological knowledge for the rational engineering and prediction of complex phenotypes is therefore one of the key challenges facing synthetic biologists and metabolic engineers today.

The previously introduced Resistome database began addressing this challenge by collating resistant *Escherichia coli* genotype and phenotype data into a single, standardized data corpus (*13, 14*). These data form the core of our effort to understand the molecular mechanisms of these phenotypes, contextualize novel mutants generated in future studies, and to develop data-driven strategies for engineering or limiting resistance. Ultimately, the goal of combining these datasets is to enable the rational construction or identification of stress-tolerant mutants by coupling increasingly inexpensive genomic data with powerful machine learning tools to build predictive models of phenotypic traits. As machine learning-guided strain engineering is increasingly an effective route for improving performance if sufficient high quality data is available (*15*–*17*), this statistical approach may eventually offer an effective route for forward engineering of phenotypes, even if understanding their underlying molecular mechanisms remains challenging (*18*).

In this work, we describe the latest improvements to the Resistome, with a particular focus on the recent development of a public user interface to enable end users to search and query the database for mutations of interest. A total for 45 additional studies have been added to the database, bringing the total to 400 curated datasets overall, and a variety of new tools to examine the implications of mutations and altered expression patterns have been included. After discussing database implementation improvements to improve accuracy and ease of use, we update our previous analysis of *E. coli* resistance at the field-wide genomic, regulatory, and proteomic levels, and then re-examine statistically significant pleiotropic interactions between phenotypes to determine the extent of pleiotropy exhibited by engineered or evolved mutants. We then evaluate the feasibility of phenotype predictions using neural networks to evaluate the Resistome as a source of training data. Finally, we discuss the challenges of data curation on a field-wide basis and possible improvements to sequencing technology that may help expand the Resistome further and enable better phenotype prediction.

## Methods and Materials

### Software Distribution and Implementation

The Resistome repository is located here: https://bitbucket.org/jdwinkler/resistome release and can be freely downloaded, modified, and redistributed so long as the original attribution is maintained. Biocyc databases must be obtained separately due to their licensing terms (*19*). Python 2.7.15 was used to implement the Python pipeline for database construction and analysis, while Python 3.6.0 was used to run Tensorflow for neural network training. The user interface is hosted on the Heroku cloud platform at https://resistome-web-interface.herokuapp.com.

### Data Sources and Database Implementation

Data were obtained by manual literature curation from 400 papers and stored in a key-value text format for later parsing and validation as previously described (*13, 14*). Phenotypes were assigned both general and specific classifications to enable efficient data analysis on subsets of the data. Each record in the Resistome corresponds to a uniquely identified study containing at least one mutant or expression dataset, along with any associated metadata. Using our custom pipeline, curated genotypes are converted into objects representing Paper, Mutant, Mutation, and G(ene)E(xpression)Change data, checked for validity, and then inserted into a Postgres 9.5 database. While currently the Resistome contains only *E. coli* data, there is no technical restriction preventing the inclusion of additional species, should the necessary reference databases be available. Depending on the analysis, Resistome data may be combined into an *E. coli* pangenome based on sequence homology between MG1655, REL606, and W genes indicated in reference databases.

Biocyc MG1655 20.0, REL606 15.5, and W 19.0 databases (*19*), converted into a custom Postgres database with an in-house script, are used to provide biological information to annotate gene properties and other host information. The database schemas, scripts needed to build the database from raw key-value data, and installation instructions are included to allow users to build their own Resistome distribution on their system.

### Supporting Tools and Data Sources

In addition to the database creation pipeline and supporting Biocyc databases, the Resistome employs a wide variety of data sources and tools to facilitate analysis of genotype-phenotype relationships. Data visualizations for this work were generated using adapted Matplotlib 2.0.2 functionality as well as Circos (*20*) for display of circular genomes. Statistical analysis was performed using Scipy 0.17.0 and Numpy 1.12.1; a maximum of *P* = 0.05 was used as the significance threshold in all tests, with more stringent significance criteria used as noted. Tensorflow 1.7.0rc1 in Python 3.6.0 was used for evaluating the practicality of phenotype prediction using neural networks.

Several conceptually similar but more targeted genotype-phenotype databases have been incorporated into the Resistome (*5*, *21*, *22*). In the case of ChemGen (*21*), only mutants with 2.5-fold or greater perturbations in growth were recorded as resistant or sensitive mutants (*13*). Estimates of the functional impact of mutations were obtained using SNAP2 and INPS (*23, 24*). *E. coli* regulatory network data was obtained from RegulonDB (*25*) and EcoliNet V1 (*26*). A recently released mapping of gene knockouts to perturbed metabolites was also added to allow users to examine mutational impacts at the metabolomic level (*27*).

### Tensorflow Model Training

Tensorflow deep neural network model training data were extracted from the Resistome in the form of sets of affected biological processes (determined by gene ontology), converted from the genotype of each mutant. For each phenotype class, these data were randomly split into training and evaluation datasets. Training data was randomly resampled with replacement to obtain 500 genotypes for training; note that some phenotype classes in the Resistome contain fewer than this number of mutants, so repeated data may contribute to overfitting for rarer genotype-phenotype combinations. After evaluating different network architectures, a single hidden layer of 32 sigmoid activation nodes was trained to predict phenotype class from biological process perturbations over 50 epochs. The network architecture and training periods were chosen empirically without extensive optimization. Each phenotype class was predicted using a distinct model; model accuracy was measured by using the evaluation dataset.

## Results and Discussion

### Database updates and quality control

Utilizing the initial release of the Resistome required users to download the distribution and other software, and then familiarize themselves with the custom software pipeline. To reduce these barriers, we initially focused on enabling users to integrate the Resistome into their own software tools. We therefore converted the Resistome into a SQL database where the same key-value data are stored as records with a well-defined structure to break this tight coupling between stored data and our software pipeline implementation. While some knowledge of a programming language and relational databases is still required to query this form of the Resistome, researchers need only the defined schema to build their own analysis tools, obviating the need to modify the provided Resistome software pipeline. Most programming languages routinely used for scientific computation are able to interface with SQL databases, allowing flexibility in the tooling used with the database.

To further simplify access, we developed a simple web interface hosted at https://resistome-web-interface.herokuapp.com that allows users to make simple gene, phenotype, or similarity-based searches of the database. Researchers can therefore perform common analyses that are needed when examining novel resistant mutants: searching for phenotypes associated with a set of mutated genes and vice versa, and identifying genotypes similar to a set of mutated or differentially expressed genes. As a result, searches can be performed once genotypic or phenotypic data is available to identify related mutations, removing the need for any manual programming and ensuring that the most up to date version of the Resistome is used.

With these accessibility enhancements in place, our focus shifted towards improving the quality of curated data. Depending on the size of a given dataset, curation fidelity can be compromised by human error by the curator if performed manually, while datasets describing hundreds or thousands of mutants are processed through custom parsers processing author-provided genetic and expression datasets. Due to the wide variability in published dataset formats and mutation annotations, the custom parsing required for large datasets is a common source of curation errors. After implementing a quality control filter to identify potential curation mistakes, we successfully detected problematic gene names, mutation locations, annotation formatting, and other errors that have since been corrected. All Resistome records conform to the minimum standards enforced by the pipeline, reducing the likelihood of observing the same errors in the future.

The second type of error is disagreement between provided literature data and current biological reference sources. For example, SNP and missense changes in the Resistome are specified using a wild-type allele or residue, location, and the resulting sequence change observed in the sequence mutants. However, 44.2% of SNP mutations in the Resistome specify wild-type alleles that do not match the corresponding base in the current *E. coli* MG1655, REL606, and W genome sequences, even after adjusting for changes in reference genomes over time. Missense mutations have a much lower rate of disagreement between curated and reference data: only 5.6% of specified wild-type amino acid residues did not match the recorded polypeptide sequence. One method for simultaneously addressing the myriad potential sources of these discrepancies and simplifying data curation may be to publish genotype data as genome difference files explicitly defining the reference genome, as seen in some studies (*28*). Alternatively, repeating genome reassembly using archived reads is an option if sufficient storage and computational resources are available, as is increasingly the case for synthetic biology laboratories.

### Revisiting the Resistome mutant collection

The Resistome currently contains 12,829 *E. coli* MG1655, REL606 (B), and W mutants with a combined total of 460 different phenotypes, accompanied by 247 gene expression datasets (Table 1). These phenotypes were not studied at random; societal needs drive examination of antibiotic resistance or solvent tolerance phenotypes within particular species due to their practical relevance to human needs. Antibiotic resistance is the most common phenotype class exhibited by Resistome mutants (Figure 1). Solvent and organic acid resistance phenotypes are also frequently characterized as part of attempts to understand product and feedstock toxicity common in industrial bioprocesses (*29, 30*). Depending on industrial and medical needs, it is possible that these trends may shift significantly in the future; for instance, identifying loci affecting phage resistance may become a research focus again due to interest in augmenting traditional antibiotic therapy with targeted phage treatments (*31*).

**Table 1:**
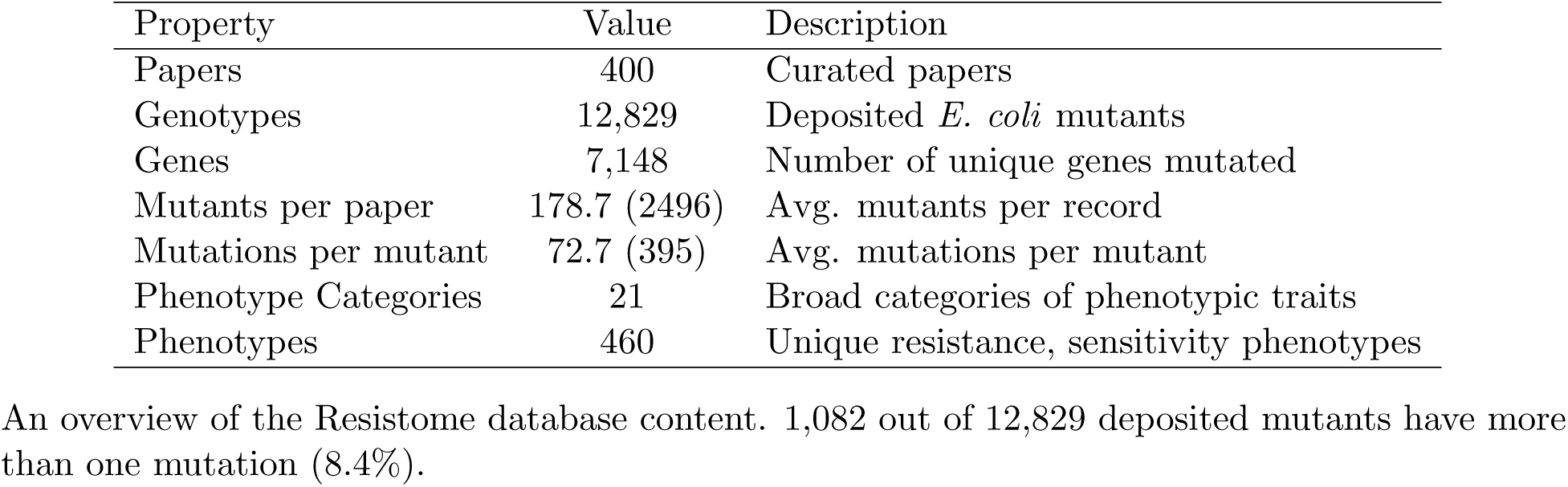
Current Resistome Properties

| Property | Value | Description |
| --- | --- | --- |
| Papers | 400 | Curated papers |
| Genotypes | 12,829 | Deposited *E. coli* mutants |
| Genes | 7,148 | Number of unique genes mutated |
| Mutants per paper | 178.7 (2496) | Avg. mutants per record |
| Mutations per mutant | 72.7 (395) | Avg. mutations per mutant |
| Phenotype Categories | 21 | Broad categories of phenotypic traits |
| Phenotypes | 460 | Unique resistance, sensitivity phenotypes |

An overview of the Resistome database content. 1,082 out of 12,829 deposited mutants have more than one mutation (8.4%).

**Figure 1:**
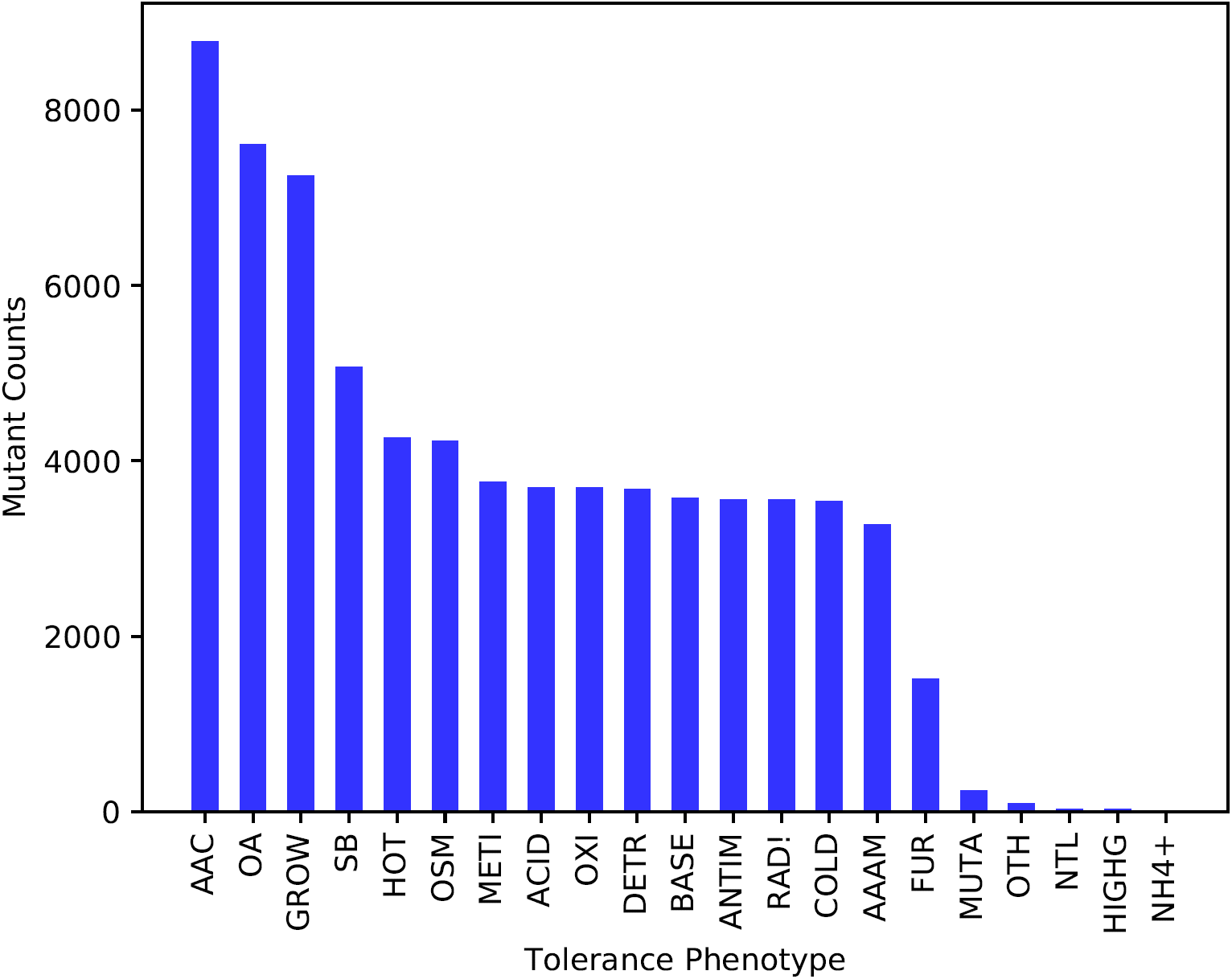
Combined counts of each phenotype category in the Resistome. Legend: AAAM: amino acid antimetabolites, AAC: antibiotics and anti-chemotherapy resistance, ACID: low pH, ANTIM: general antimetabolites, COLD: low temperature, DETR: detergents, FUR: furfural, GROW: growth in the absence of a specific stressor, HIGHG: high gravity, HOT: high temperatures, METI: metabolic inhibitors, MUTA: mutagens, NH4+: quarternary ammonium compounds, NTL: nutrient limitation, OA: organic acid, OSM: hypo- or hyperosmotic stress, OXI: oxidative stress, RAD!: ionizing radiation, SB: solvents and biofuels.

The scale of genotype-phenotype interrogation continues to grow, as each curated study contains 179 mutants on average. However, 92% of all mutants, being principally derived from genome-scale libraries, possess a single mutation per strain. The remaining mutants, with an average of 73 mutations per strain (*σ* = 395), were obtained from studies using random mutagenesis or adaptive evolution that routinely generate lineages of mutants with multiple mutations due to their large population sizes and mutation rates (*32, 33*). Some of the newly curated studies also contain mutator strains, leading to an inflation in the mutations per strain statistics compared to the original Resistome release. While untargeted and prone to accumulating hitchhiking mutations, random selections are able to explore combinations of mutations that are otherwise inaccessible for designed libraries due to the combinatorial explosion of mutation possibilities: *E. coli* MG1655 has 4,501 genes, so a library containing every double mutant has on the order of nine million mutants if synthetic lethality is neglected. While still an infeasible scale, parallel editing technologies that could enable less targeted multi-locus genomic libraries continue to improve rapidly (*34*–*37*).

The differences between random and rationally-designed library studies are similar to those found previously (Table 2). Overall, a few mutation types in the Resistome predominate (Figure 2): deletions, transposon insertions, and SNPs comprise the majority of observed mutations, with designed mutants containing mainly deletion and overexpression mutations compared to the 10.8 different types of mutations in random mutants on average. Mutations affecting essential genes are underrepresented in targeted libraries as a result of their limited mutational spectra, as essential genes are mutated 10.5 times more often in random selections. These differences in the constraints governing random and designed library studies translate into more comprehensive perturbation of gene ontology biological processes for random studies (36.9 vs. 9.5 GO biological processes, *P* = 1.7 * 10^-278^, two-tailed t-test),which may indicate generating diverse mutations within a host may be more effective at identifying genotype-phenotype relationships than gene deletion or overexpression only.

**Table 2:** Comparison of Random and Designed Mutants

| Property | Random | Designed | P-value |
| --- | --- | --- | --- |
| Mutation types | 10.81 | 1.33 | $2.58 * 10^{-100}$ |
| Mutations | 9.29 | 1.04 | $3.46 * 10^{-90}$ |
| GO perturbations | 36.87 | 9.52 | 0 |
| Essential genes modified | 0.67 | 0.063 | 0 |

A comparison of random mutants derived from adaptive laboratory evolution or other types of selection experiments versus designed, generally library-derived strains. Statistical comparisons were performed using two-tailed Students t-tests.

**Figure 2:**
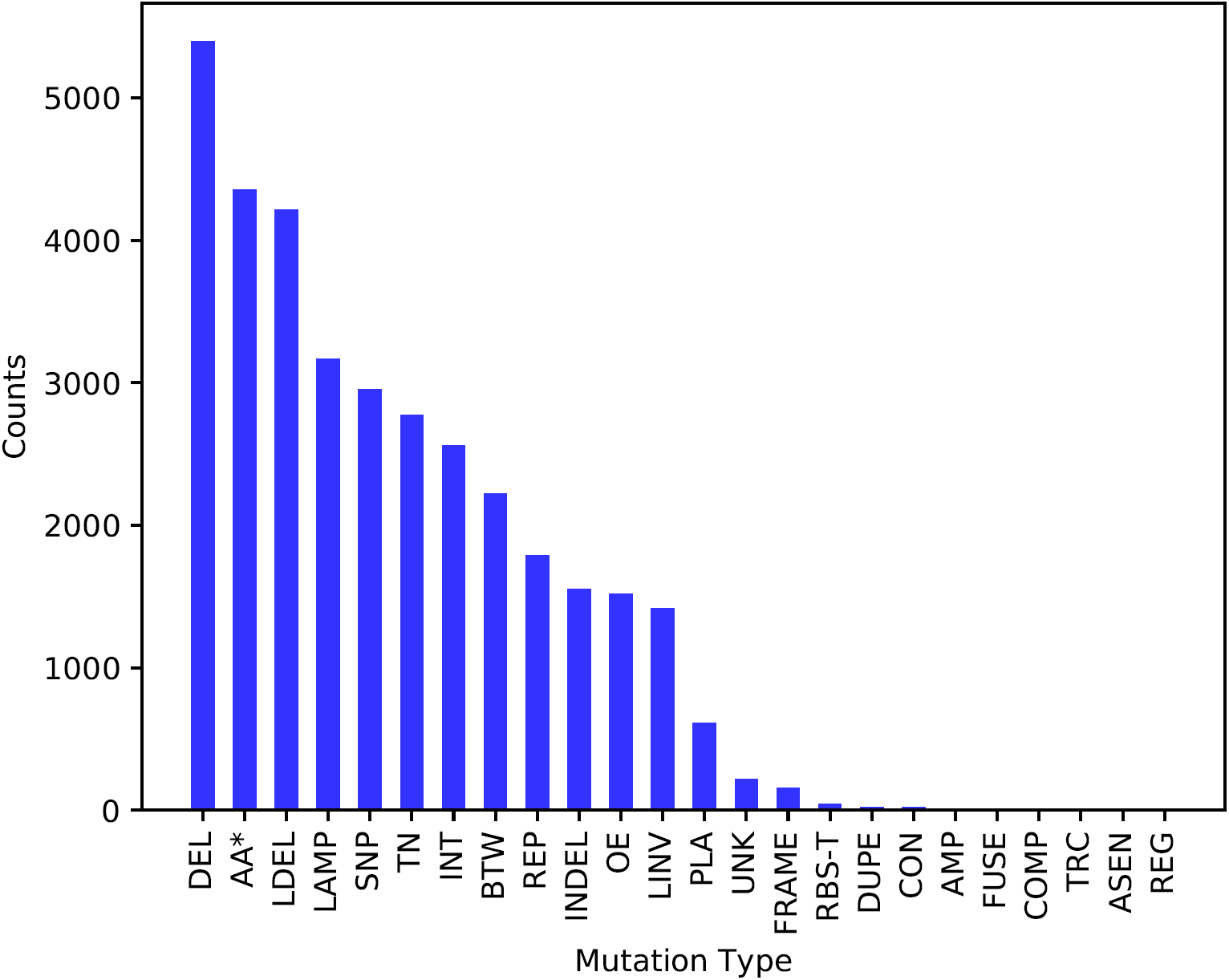
Counts of each mutation type listed in Resistome mutants. The full names for each type are: AA*: missense mutation, AMP: amplification, ASEN: antisense gene knock-down, BTW: intergenic mutation, COMP: compartmentalization, CON: constitutive expression, DEL: deletion, DUPE: internal duplication, FRAME: frameshift, FUSE: protein fusion, INDEL: small (¡100 bp) insertion or deletion, INT: genomic integration, LAMP: large amplification, LDEL: large deletion, OE: overexpression, PLA: plasmid cloning, RBS-T: RBS tuning, REG: new regulator relationship, REP: gene repression, SNP: single nucleotide poly-morphism, TER: terminated, TN: transposon insertion, TRC: truncation, UNK: unknown. Note that residue changes and SNPs included in the database represent distinct mutations, and both categories can include silent mutations. Mutation Separation (kilobases)

### An overview of Resistome mutations

There are approximately 29,000 mutations in 7,148 unique genes over all three *E. coli* strains recorded in the Resistome. A complete list of mutated genes, including their associated phenotypic and annotation metadata, is available in Table S1. Mutational coverage of the genome (Figure 7) is almost complete: only a small fraction (176/4501, 3.9%) of MG1655 genes are not yet associated with at least one resistance phenotype, speaking to the effectiveness of genomic libraries, random selections, and higher throughput screening to identify genotype-phenotype linkages. Non-mutated genes are generally transfer RNA or transposon-related based on their annotated functions (Table 3); their apparent lack of phenotypic impact is probably due to a combination of these genes genuinely not affecting screened phenotypes directly and not being included in most genomic libraries. The 25 most frequently mutated genes account for 6.7% of all mutated genes, about 19.3-fold more than would be expected than the null expectation that all genes are equally likely to influence resistance.

**Table 3:** Ontological properties of non-mutated genes

| GO Tag | Description | P-value |
| --- | --- | --- |
| GO:0006412 | Translation | $9.93 * 10^{-6}$ |
| GO:0012501 | Programmed cell death | $2.99 * 10^{-6}$ |
| GO:0022625 | Cytosolic large ribosomal subunit | $7.74 * 10^{-7}$ |
| GO:0006310 | DNA recombination | $3.62 * 10^{-12}$ |
| GO:0004803 | Transposase activity | $3.48 * 10^{-15}$ |
| GO:0005737 | Cytoplasm | $5.80 * 10^{-16}$ |
| GO:0006313 | Transposition, DNA-mediated | $1.16 * 10^{-16}$ |
| GO:0032196 | Transposition | $6.33 * 10^{-18}$ |
| GO:0005829 | Cytosol | $1.15 * 10^{-21}$ |
| GO:0030533 | Triplet codon-amino acid adaptor activity | $1.49 * 10^{-65}$ |

Gene ontology tags that are enriched for non-mutated genes compared to the background *E. coli* genome. Statistical significance was assessed using Fishers exact test.

Mutated genes related to transcription, translation, DNA replication, and maintaining ion homeostasis are frequently enriched compared to that expected for the *E. coli* genome. Each class has peculiarities that account for differences in stress conditions; for example, an increase in mutations affecting topoisomerase functions is observed for antibiotic (fluoroquinolone) resistance. A complete list of these enriched tags and their significance is provided for reference in Table S2. While these findings are not surprising overall, they serve as a reference for researchers to understand which biological processes are likely to be affected if they are attempting to generate novel mutants.

Pleiotropy appears to be common among resistant mutants due to the number of phenotypes associated with each gene (Figure 3A) across the database. Regulator-driven adaptation is an excellent example of this phenomenon, as mutated regulators frequently confer resistance phenotypes across the entire range of phenotype classes in the Resistome (Figure 8). Since understanding the root cause of phenotypes associated with the mutation of global regulators typically requires extensive analysis following their identification (*38*–*41*), finding routes to limit the mutation of regulators and other genes with outsized phenotypic impacts may, paradoxically, simplify the characterization and subsequent re-engineering of resistance-conferring mutations into other strain backgrounds. After counting gene-phenotype mappings identified in the Resistome, only 326 MG1655 genes were associated with one resistance phenotype. One to one mappings between mutations and a phenotype appear to be rare, implying that most mutants will need to be re-engineered or selected with care to reduce off-target traits associated with evolved biocatalysts (*42*).

**Figure 3:**
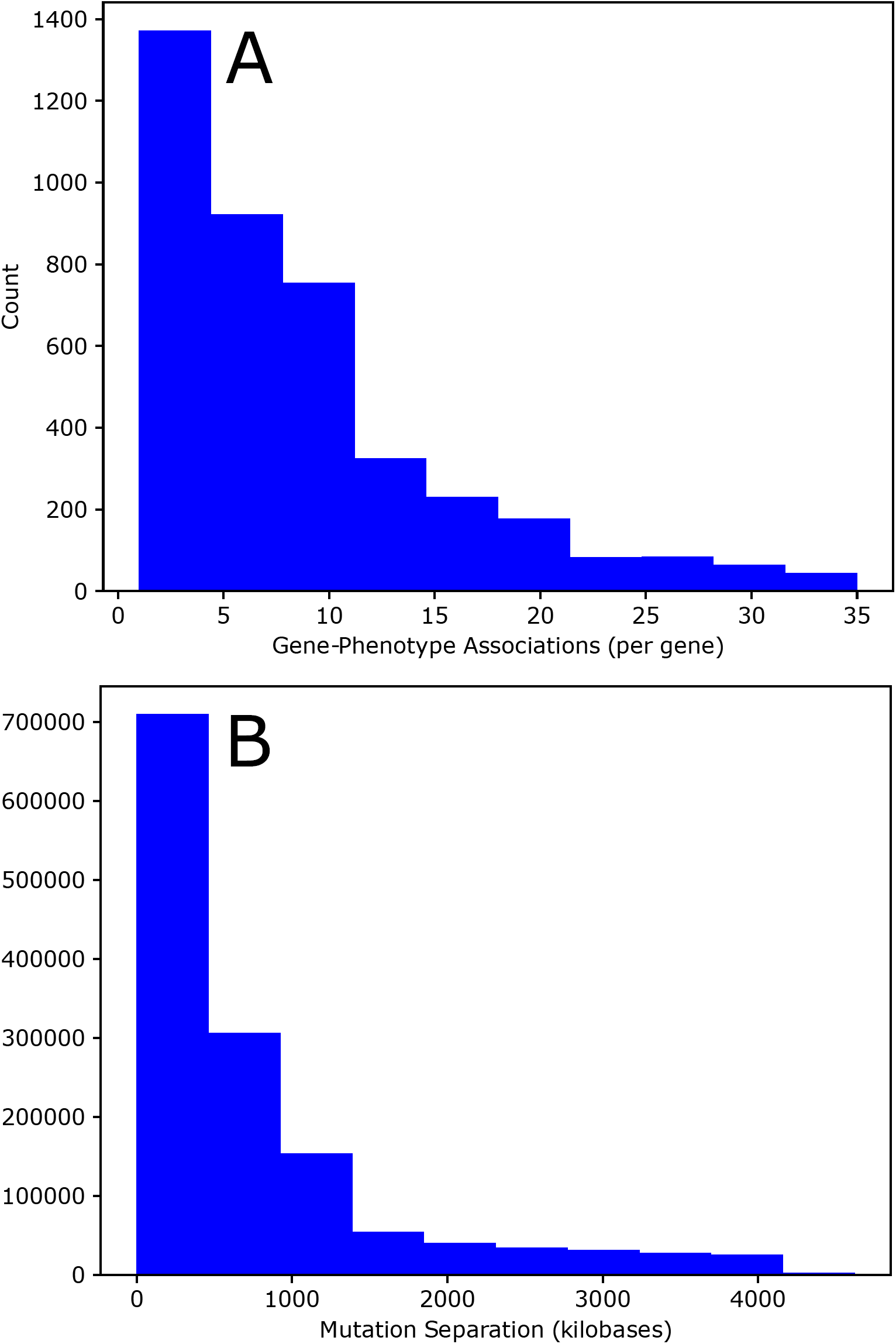
A. Distribution of the number of phenotypes associated with a single gene extracted from Resistome mutants. B. Distribution of distances between all pairs of mutations occurring in the same genotype for the 1,082 mutants with more than a single mutation. Note that the total number of unique mutation pairs within each strain is equal to *N* ^2^*/*2, where *N* is the number of mutations.

Sequencing technology plays a key role in rapid genotype interrogation for all types of resistance-related studies. Assigning mutations within duplicated regions and genotyping mixed populations of unknown genotypes remains challenging as mutations are separated by a median of 440 kilobases (Figure 3B). Clonal isolates must therefore be obtained for genotyping as otherwise short reads cannot provide linkage information, though the frequencies of individual mutations within the population can still be tracked (*32*, *43*, *44*). Designed genome-scale libraries circumvent this issue with genotype-associated barcodes that can fit into a single read, enabling the use of population-level sequencing to assign fitness scores without handling isolates (*10, 45*). A combination of long-read sequencing technology to capture multiple mutations per read and algorithmic improvements to reconstruct the corresponding complete genotypes may therefore allow faster characterization of novel genotype-phenotype relations, as has already been demonstrated elsewhere for defined mixed populations (*46*).

### 0.1 Genomic and proteomic mutation signatures

Mutations are not randomly distributed throughout the genome, independent of sequence content and context, and the resulting mutational biases have a significant impact on resistance acquisition. Indels, one of the major classes of mutations to occur randomly in evolved mutants, often occur in or near homopolymeric runs (contiguous identical bases) as a result of polymerase slippage (*47*) and can disrupt coding sequences and regulatory sites depending on where they occur. For the 1,390 insertions or deletions included in the database that can be localized to the MG1655 genome, 26.7% occur in the vicinity of (not necessarily within) a homopolymeric run containing at least three identical bases. The majority (1142/1390) of these indels are found outside of coding sequences.

SNPs are the second type of mutation to occur frequently in Resistome mutants, and can be recorded as explicit nucleotide mutations or indirectly as residue changes within polypeptides; in the latter case, the likely underlying nucleotide change can often be inferred from genetic code constraints on amino acid replacements (*48*). *E. coli* SNPs are known to be biased towards transition (G/C to A/T) mutations in the absence of selection due to differential repair efficiencies (*47*). G to A and C to T mutations are the most frequently observed explicit or inferred SNPs, accounting for approximately 28% of all changes (Figure 4). There are some differences in the relative abundance of nucleotide substitutions for genomic versus coding sequence changes; comparing explicitly annotated SNPs, including both SNPs found in coding and non-coding regions, to those inferred from amino acid changes reveals that G to C, C to G, T to A, and to a lesser extent, T to G mutations are slightly overrepresented in mutations within coding sequences but exhibit the same overall bias towards transition mutations (data not shown).

**Figure 4:**
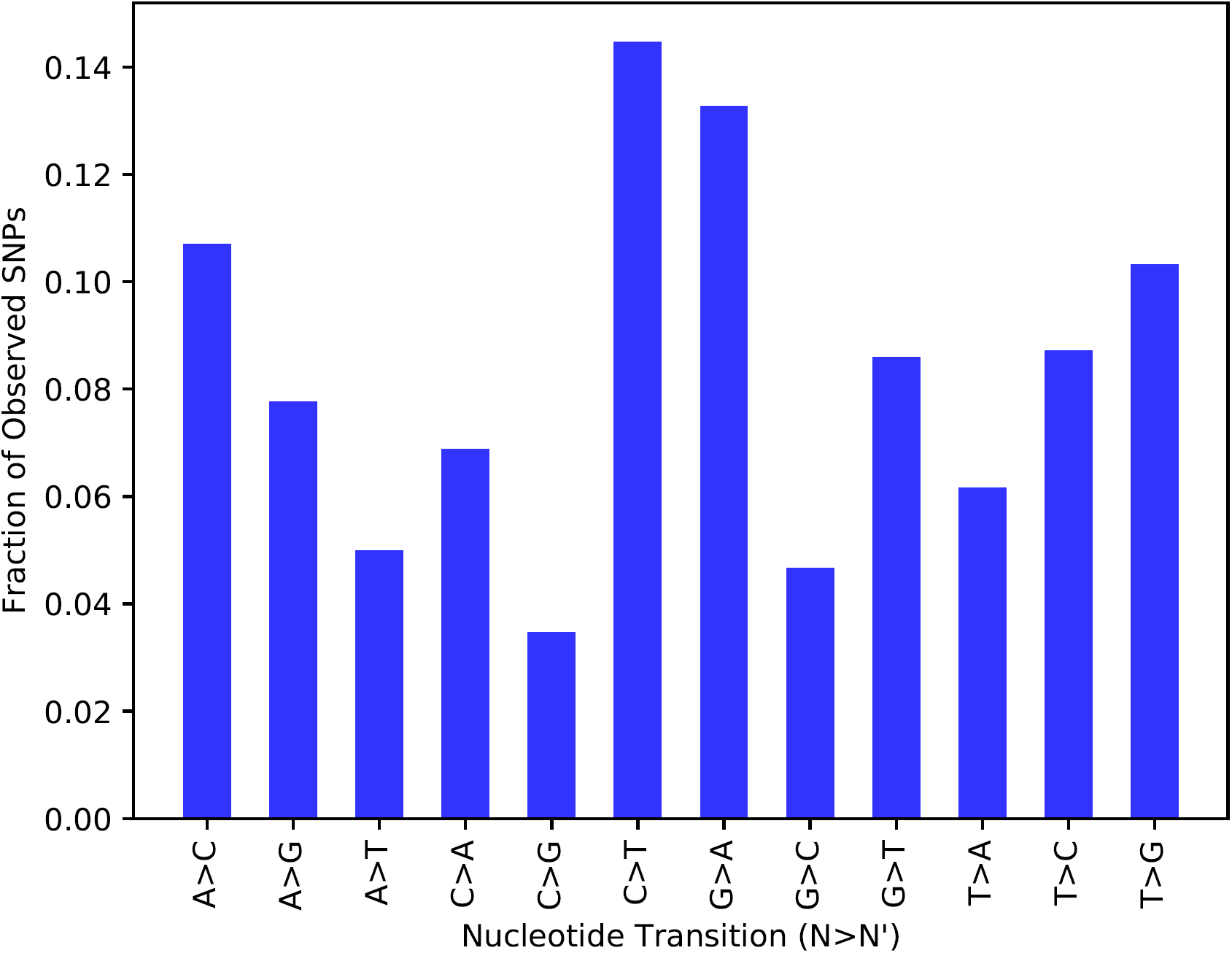
Observed nucleotide base replacements for all Resistome SNPs, assuming all base changes are specified correctly despite ambiguity concerning their genomic location. Residue replacements are included if the underlying codon change can be deduced from genetic code constraints.

When examining mutations affecting codon sequences, the possible residue changes are only sparsely sampled in the curated mutational data (Figure 5A). The majority (91.1%) of these amino acid changes require only a single SNP, as shown in Figure 5B. The frequency of residue to residue changes is significantly inversely correlated with the number of SNPs required to convert between their underlying codons (Spearman *R* = *−*0.64, *P* = 6.8*\**10^-52^). Experiments relying on natural mutation mechanisms or error-prone PCR are particularly impacted by this effect, given the rarity of multiple nucleotide changes within the same codon and the natural error buffering of the genetic code (*48*). High throughput genome engineering using synthesized DNA followed by selection (*9*, *34*, *49*) is being used on a larger scale to circumvent this limitation, as a considerable number of annotated residue replacements now require a minimum of two or three nucleotide changes. These approaches may therefore improve the accessibility of high fitness genotypes and the effectiveness of saturation mutagenesis (*50*).

**Figure 5:**
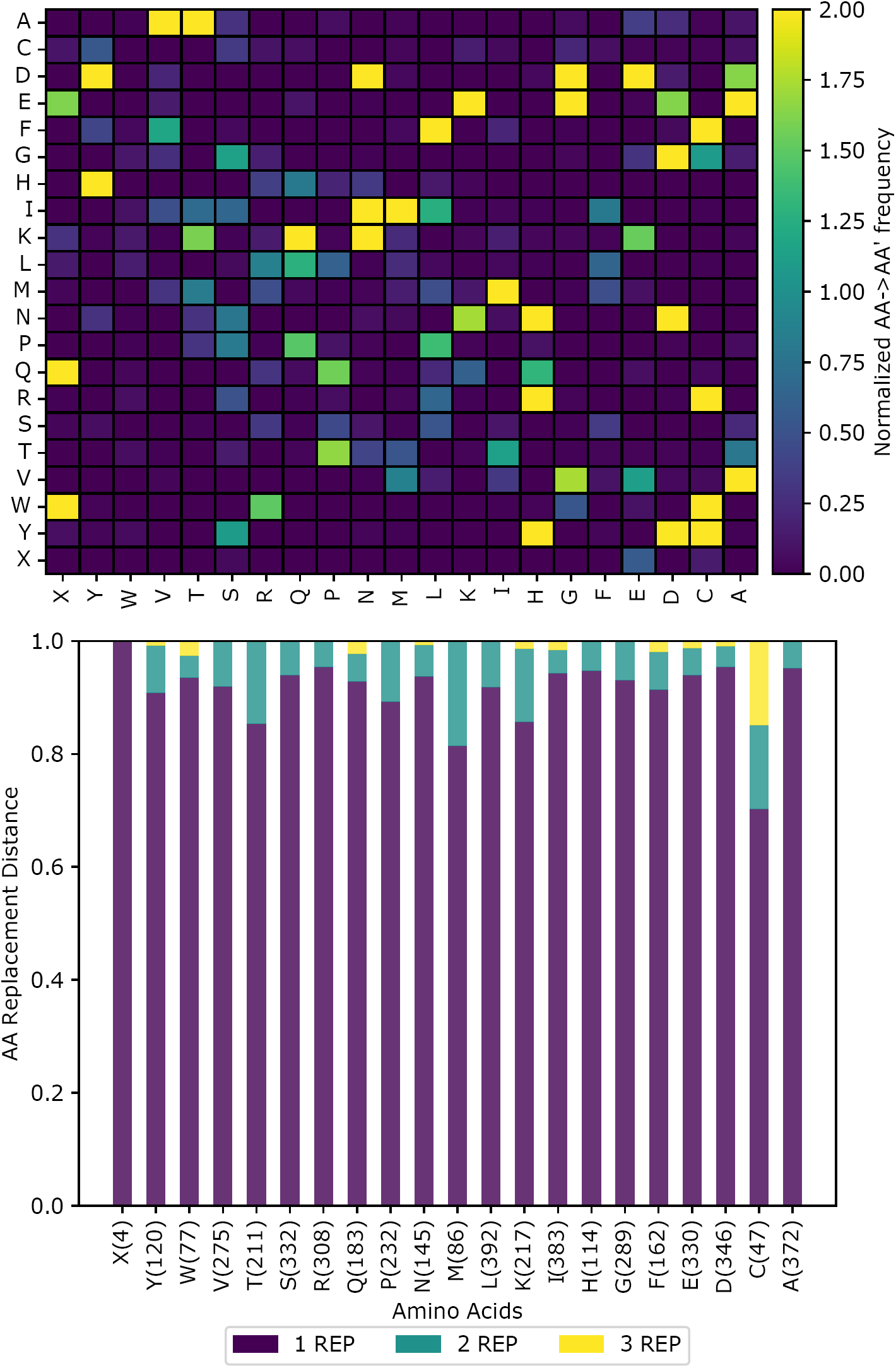
Top: Map of observed residue interconversion within all Resistome mutants, normalized by the number of unique gene-mutation pairs occurring over the database multiplied by the number of codon-codon pairings possible between each amino acid. Bottom: Proportion of amino acid changes requiring one, two, or three nucleotide changes for each observed mutation, along with the count of how many times a residue change results in each amino acid in parenthesis.

Predicted changes in free energy associated with recorded residue changes suggests that the majority of mutant proteins are slightly more stable than their original polypeptides with the median ddG being −0.49 kcal/mol using the INPS method (*24*). Since protein stability is evolutionarily fine-tuned to ensure proper folding but with sufficient flexibility to accomplish substrate binding or catalysis (*51*), these data imply some degree of functional disruption for the variant proteins, especially given the statistically significant overrepresentation of missense mutations in DNA-binding motifs (4.28-fold overrepresentation by Fishers exact test, *P* = 1.24 *\** 10^*-28*^). Additional study is needed to better understand the precise impact of these coding sequence changes before firm conclusions can be drawn about their cellular impact. The Resistome also provides a tool to combine the predicted effect of amino acid replacement, calculated using SNAP2 (*23*), with a visualization of recorded mutations within a protein sequence of interest (e.g. Figure 6) to aid researchers in selecting additional sites for mutagenesis.

**Figure 6:**
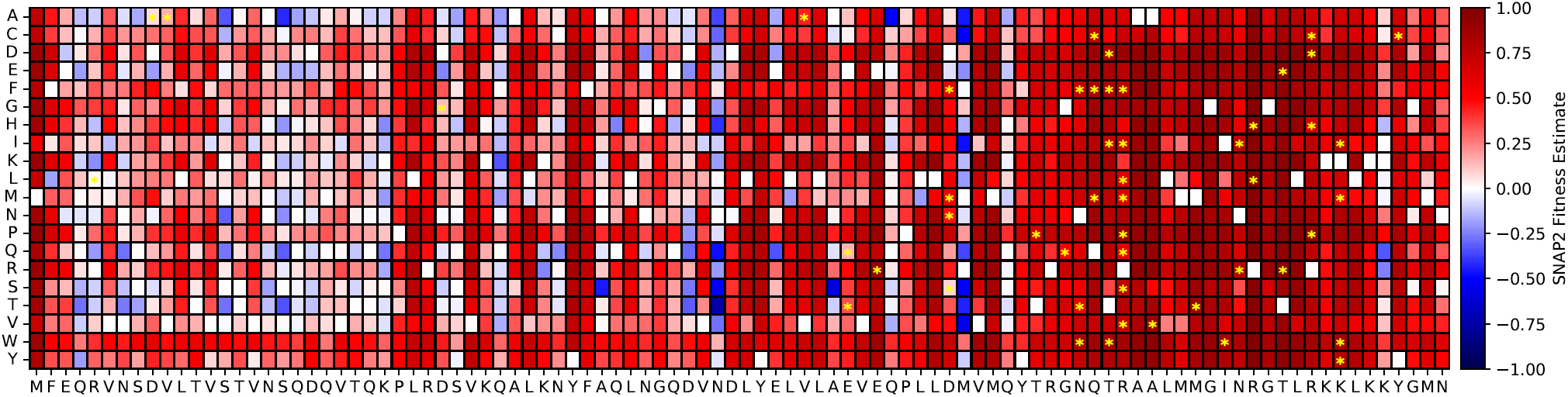
Combining functional predictions for amino acid replacement obtained from SNAP2 with deposited mutations for B3261 (*fis*). White boxes indicate the wild-type residue in the original sequence, and yellow stars indicate that a given mutant residue occurs in a Resistome mutant. Potential replacement residues on the leftward y-axis are colored according to their predicted effect on protein function; the more negative the score, the stronger the prediction of replacement neutrality. Conversely, more positive scores indicate a higher predicted likelihood of the mutation impacting protein function.

### Evaluating phenotype interactions and predicting traits

Based on our analysis of resistant mutants so far, weaving together a cohesive, mechanistic understanding of how a set of observed genotypes creates the linked phenotypes remains a daunting challenge. Some phenotypes, particularly those involving antibiotic resistance, have well-defined targets and resistance mechanisms (*22*); others affect a wider array of cellular components and accordingly their genotypes are harder to predict (*4*, *52*, *53*). This complexity fundamentally arises from the mutations themselves, interactions with host background and co-occurring mutations, and peculiarities associated with the environment used for selection that may be undocumented or unknown altogether. Mutants evolved for complex phenotypes often exhibit pleiotropy as a consequence of these factors and may not be reproducible even when ostensibly repeated under nominally identical conditions (*7*).

While it may be impractical to accurately predict the phenotypes or genotypes of specific resistant mutants, having statistical evidence of a relationship between phenotype classes may allow researchers to better target their strain characterization efforts once resistant mutants have been identified. One route to assess the possibility of an interaction is to evaluate the significance of the number of shared genes between phenotype category pairs using Fishers exact test for an *E. col* i-sized genome, as shown in Figure 9). While this approach does not account for the actual biological processes perturbed, it nonetheless provides additional support for the pervasiveness of pleiotropy among resistant mutants. Collateral antibiotic resistance, in particular, may be a side effect of phenotypic improvement due to overlaps with oxidative, thermal, and acid resistance. Definitive assessment of collateral phenotypic interactions will require screening mutants against a battery of stress conditions to determine their complete set of resistance traits, as it is rare that adaptive mutants are comprehensively screened to determine their traits, with some exceptions (*21*).

**Figure 7:**
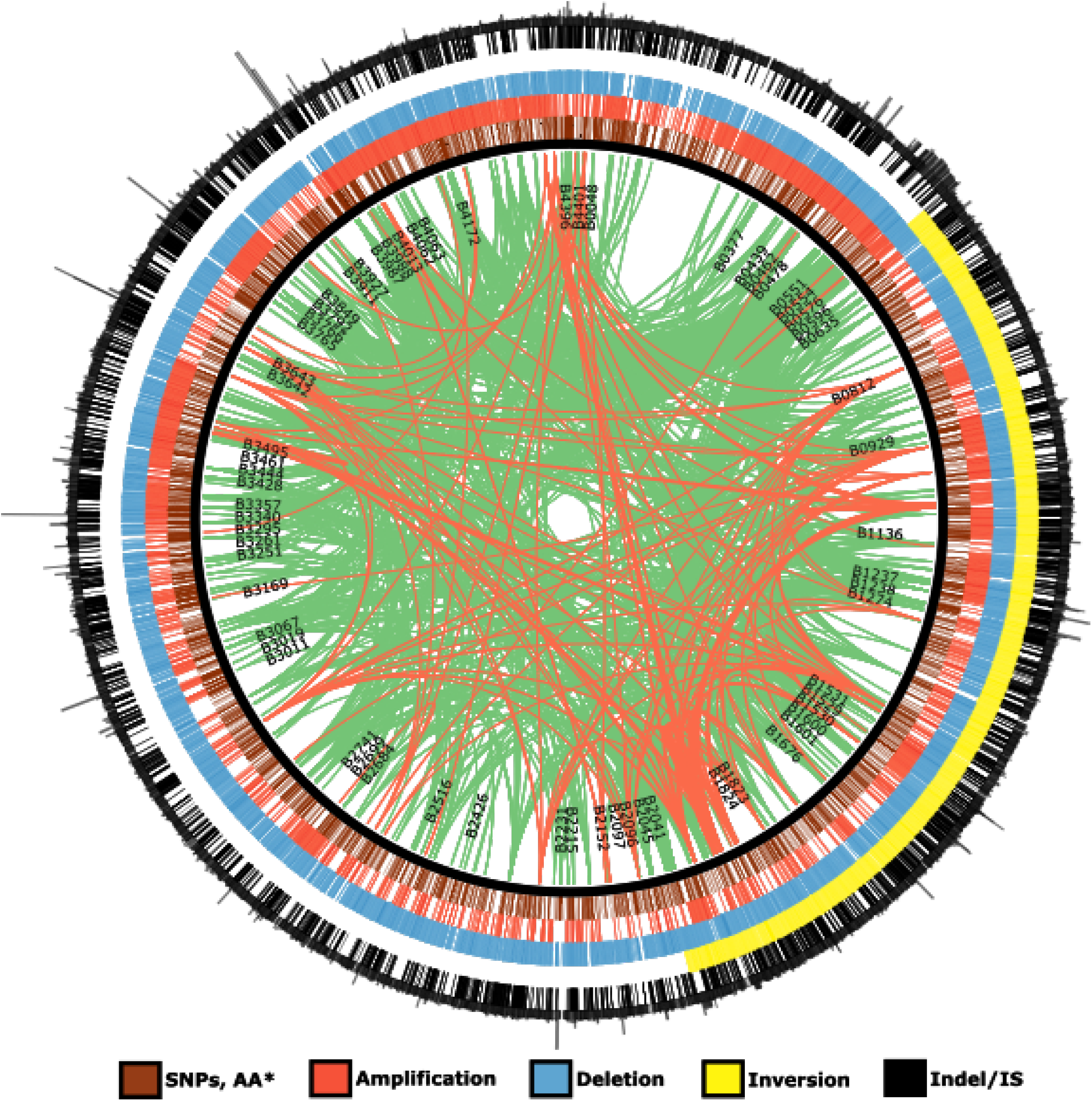
Visualization of the current *E. coli* mutome recorded in the Resistome; mutation types are denoted by position (SNPs and missense mutations, amplifications, deletions, inversions, and finally indels) and color. Spikes are proportional to mutation frequency.

**Figure 8:**
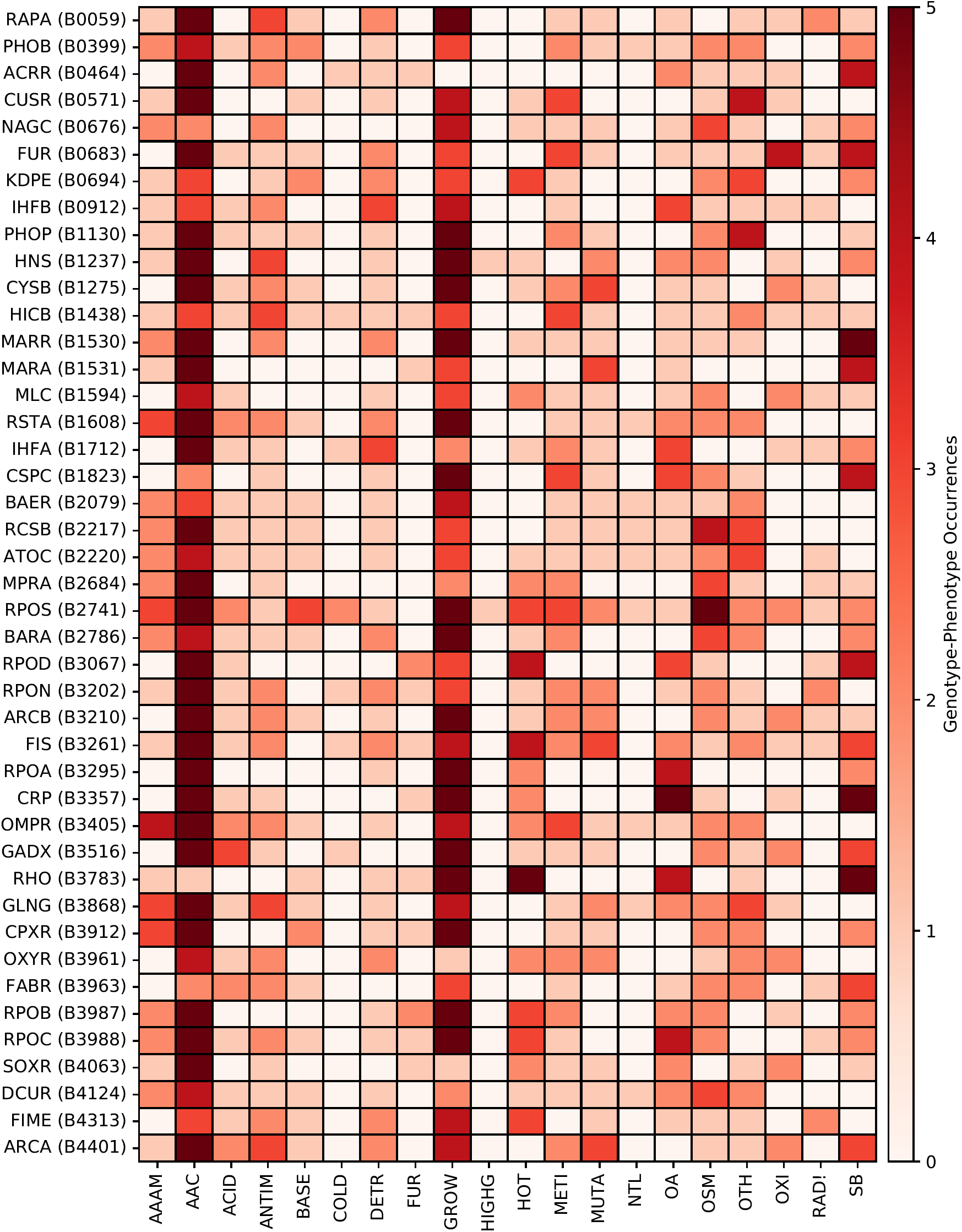
Map of how often each regulator (Y axis) is associated with a phenotype in each category (X axis). Each regulator listed is associated with at least twenty phenotypes total. Legend entries are the same as Figure 1.

**Figure 9:**
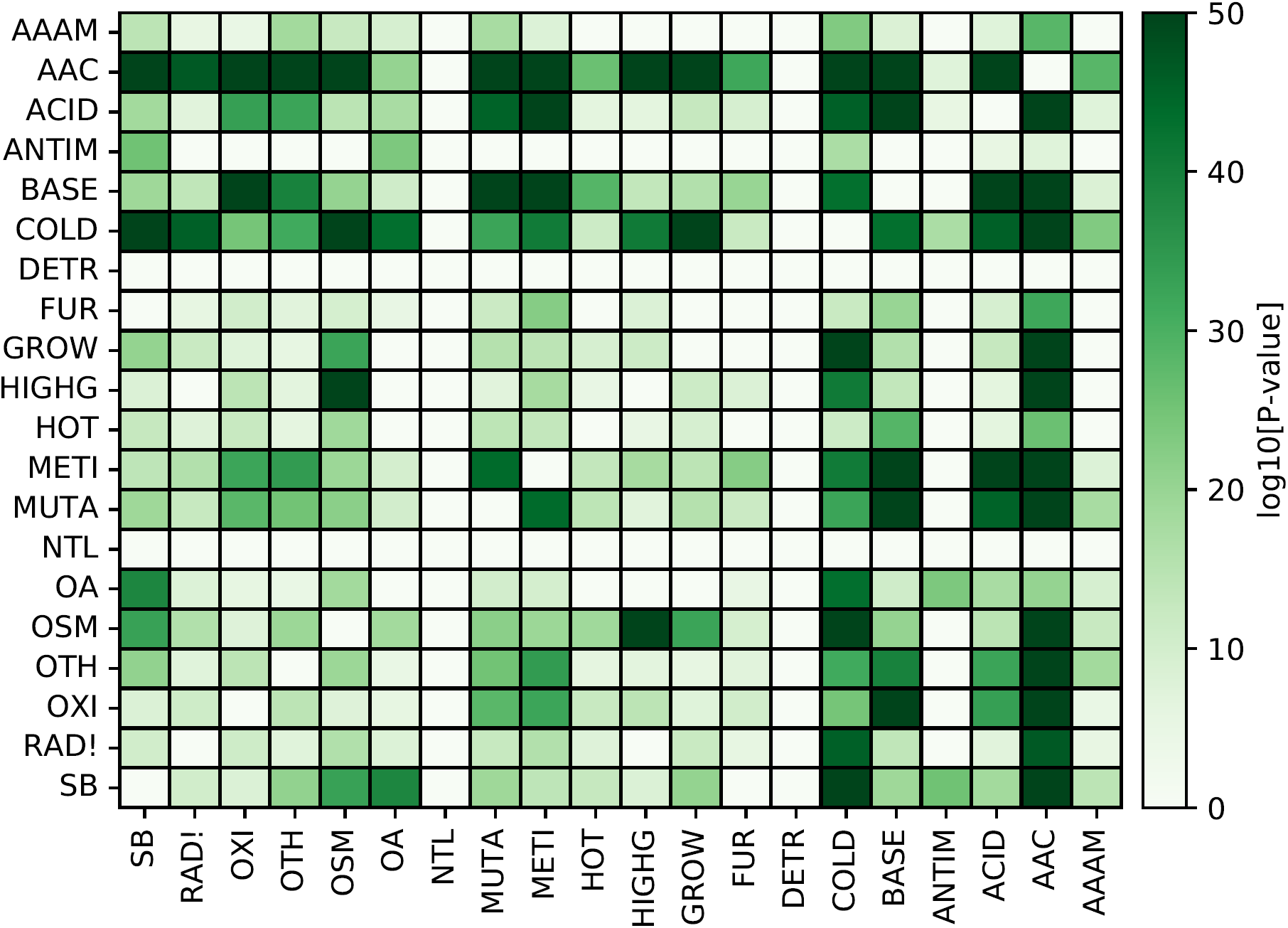
Statistically significant enrichment or depletion of shared genes for each phenotype category. P-values are capped at 10^−50^ for visual clarity. Legend entries are the same as Figure 1.

Inhibitors do not impose a single type of stress on an organism; rather, each mutant genotype triggers a constellation of cellular changes to yield the corresponding phenotypes in each mutant. While rendering mechanistic prediction of phenotype from genotype is challenging, statistical models based on neural networks offer an alternative for the prediction of phenotypes from genotype datasets (*54*). Our attempts to train neural networks to predict phenotype class from the genotype generally resulted in prediction accuracies on the order of 55-70% (Figure 10A). This level of accuracy is low compared to that expected from most neural network applications with more training data (*55*), and the correlations between phenotype class predictions presented in Figure 10B are generally negligible despite there being known correlations between some phenotypes (*56, 57*). Model overfitting due to the small size of the Resistome and data resampling is likely the cause of these issues, as it would result in models that generalize poorly to new genotypic data; for comparison, DCell, a neural network-based approach for predicting growth rate from *Saccharomyces cerevisiae* genotype, relied on a training dataset containing up to 8 million standardized genotype-growth rate measurements (*54, 58*), in contrast to the approximately 13,000 Resistome mutants with heterogeneous fitness metadata. Developing predictive and generalizable phenotypic models along the lines of the used for DCell will probably require additional genotype-phenotype data with quantitative fitness scores, generated using high throughput resistance screening.

**Figure 10:**
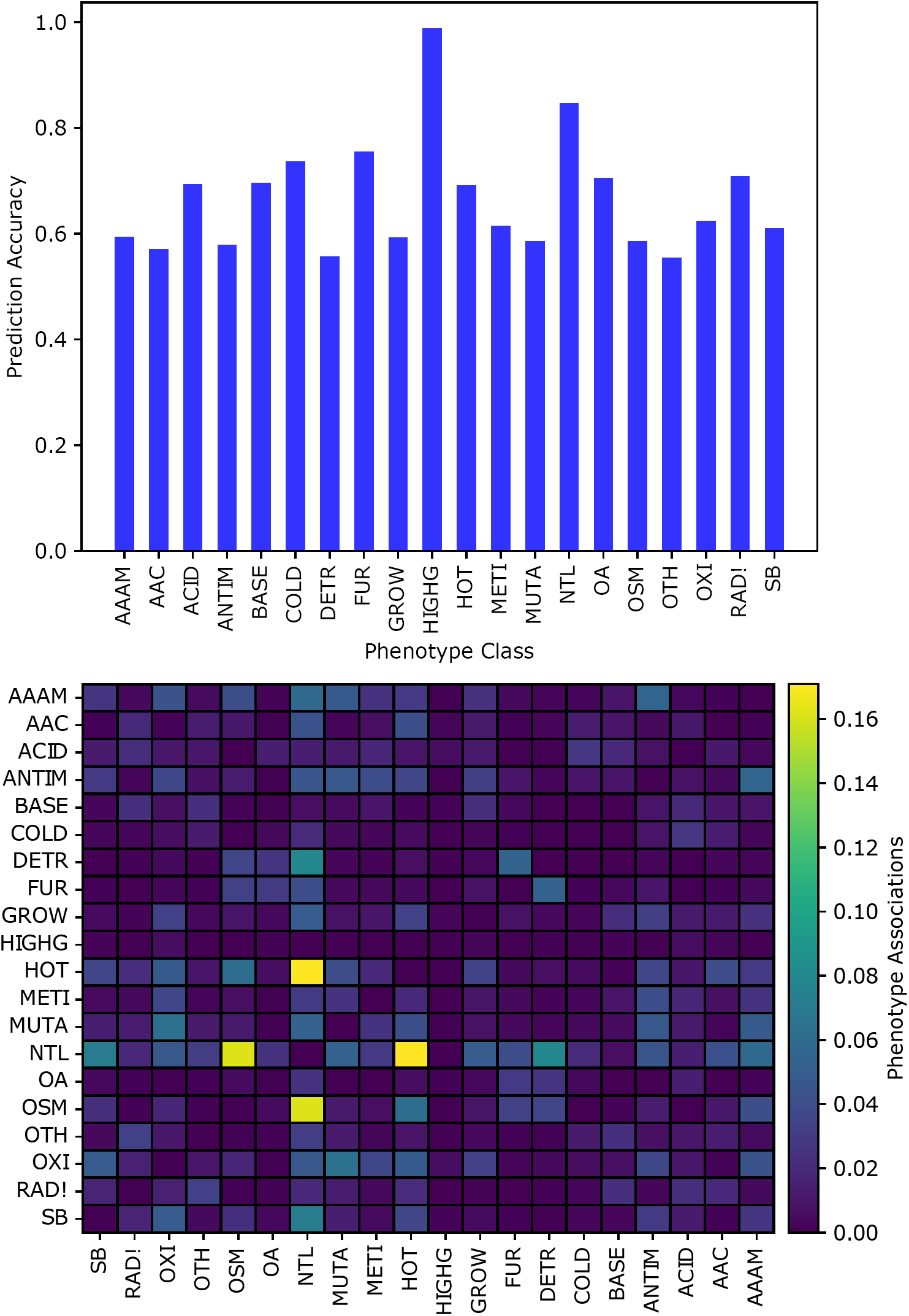
Top: Phenotype prediction accuracy using Tensorflow. Legend entries are the same as Figure 1. Bottom: Spearman R-values between model predictions for different phenotype categories; the diagonal representing self-similarity was set to zero for all phenotype categories.

## Conclusions

This work marks the release of a significant enhancement of the Resistome, both in terms of additional genomic and transcriptomic data added and a web interface to simplify access to the database. After reviewing the properties of curated mutants and biological processes affecting phenotype expression at a high level using this expanded corpus, the genomic, proteomic, and regulatory signatures of resistance were systematically explored to better define how our ability to improve phenotypes may be impacted by the underlying biological constraints at each level of cellular organization. Phenotype interactions viewed through the lens of genes shared between mutants exhibiting distinct resistance traits supports our original conclusion that pleiotropy is endemic among resistance mutants, and more extensive phenotypic screening is needed to identify collateral phenotypes when engineering more robust industrial biocatalysts or characterizing medically-relevant isolates. Attempts at phenotype prediction using neural networks were unable to reproduce this inferred pleiotropy, and the trained models appeared to overfit the small training dataset available and did not capture known phenotypic overlaps between resistance categories.

Several potential routes for addressing these limitations through database expansion were identified in the course of this study. Adopting a standard for disseminating *E. coli* genotypes would be helpful for future curation efforts and contextualizing data as updates are made to biological databases over time, and could be as simple as ensuring raw data is deposited into the NCBI sequence read archive and distributing genome difference files generated using tools like breseq (*59*) as supplementary datasets. Being able to identify complete genotypes using long-read sequencing technology within mixed populations of unknown composition would also enable rapid identification of epistatic interactions between distant mutations without requiring low-throughput single isolate analysis. Given the small fraction of Resistome mutants with multiple mutations, this step appears to be rate-limiting compared to genome-scale libraries were dynamics can be tracked by easily sequenced barcodes. At the same time, genome engineering technologies leveraging DNA synthesis, CRISPR-Cas9 genome editing, and barcoding are particularly promising for *in vivo* multiplexed random mutagenesis that may offer an alternative to purely random selections that are easily interrogated using common sequencing methodologies.

Future expansion of the database will focus on incorporating larger-scale studies generated using these advancements in genetic engineering and omics technologies, including metabolomics and proteomics datasets as they become more commonly available (*17, 60*). Regardless of the specific advances in synthetic biology, it seems certain that our abilities to engineer mutants and interrogate the biological underpinnings of their traits will continue to expand rapidly. Much like the revolution currently underway in machine learning-guided industrial metabolic engineering, flexibly collecting and combining these data streams using tools such as the Resistome with advances in machine learning offers one potential route to achieve the long-sought after goal of predicting and rationally engineering complex phenotypes.

## Acknowledgement

We thank Emily Freed and Gur Pines for their insightful comments during the preparation of this manuscript. Several researchers or research groups also kindly supplied raw data for curation, including Joan Slonczewski, Jeffrey Barrick, Brandon Gaut, Andrea Gonzlez, Rongming Liu, and Andrew Garst, while Castrense Savojardo and Jonas Reeb, respectively, provided INPS and SNAP2 missense effect predictions for Resistome mutation data.

## References

1. Blair, J. M., Webber, M. A., Baylay, A. J., Ogbolu, D. O., and Piddock, L. J. (2015) Molecular mechanisms of antibiotic resistance. Nature Reviews Microbiology 13, 42–51.

2. Huang, M., Peabody, G., and Kao, K. C. Metabolic Engineering for Bioprocess Commercialization; Springer, 2016; pp 73–100.

3. Mukhopadhyay, A. (2015) Tolerance engineering in bacteria for the production of advanced biofuels and chemicals. Trends in Microbiology 23, 498–508.

4. Dragosits, M., Mozhayskiy, V., Quinones-Soto, S., Park, J., and Tagkopoulos, I. (2013) Evolutionary potential, cross-stress behavior and the genetic basis of acquired stress resistance in *Escherichia coli*. Molecular Systems Biology 9, 643.

5. Nyerges, á. et al. (2018) Directed evolution of multiple genomic loci allows the prediction of antibiotic resistance. PNAS

6. Wytock, T. P., Fiebig, A., Willett, J. W., Herrou, J., Fergin, A., Motter, A. E., and Crosson, S. (2018) Experimental evolution of diverse *Escherichia coli* metabolic mutants identifies genetic loci for convergent adaptation of growth rate. PLoS Genetics 14, e1007284.

7. Kram, K. E., Geiger, C., Ismail, W. M., Lee, H., Tang, H., Foster, P. L., and Finkel, S. E. (2017) Adaptation of *Escherichia coli* to long-term serial passage in complex medium: evidence of parallel evolution. MSystems 2, e00192–16.

8. Jahn, L. J., Munck, C., Ellabaan, M. M., and Sommer, M. O. (2017) Adaptive laboratory evolution of antibiotic resistance using different selection regimes lead to similar phenotypes and genotypes. Frontiers in Microbiology 8, 816.

9. Liu, R., Liang, L., Garst, A. D., Choudhury, A., i Nogué, V. S., Beckham, G. T., and Gill, R. T. (2018) Directed combinatorial mutagenesis of *Escherichia coli* for complex phenotype engineering. Metabolic Engineering 47, 10–20.

10. Garst, A. D., Bassalo, M. C., Pines, G., Lynch, S. A., Halweg-Edwards, A. L., Liu, R., Liang, L., Wang, Z., Zeitoun, R., Alexander, W. G., and Gill, R. T. (2017) Genome-wide mapping of mutations at single-nucleotide resolution for protein, metabolic and genome engineering. Nature Biotechnology 35, 48.

11. Baba, T., Ara, T., Hasegawa, M., Takai, Y., Okumura, Y., Baba, M., Datsenko, K. A., Tomita, M., Wanner, B. L., and Mori, H. (2006) Construction of *Escherichia coli* K-12 in-frame, single-gene knockout mutants: the Keio collection. Molecular Systems Biology 2.

12. Kitagawa, M., Ara, T., Arifuzzaman, M., Ioka-Nakamichi, T., Inamoto, E., Toyonaga, H., and Mori, H. (2006) Complete set of ORF clones of Escherichia coli ASKA library (a complete set of E. coli K-12 ORF archive): unique resources for biological research. DNA research 12, 291–299.

13. Winkler, J. D., Halweg-Edwards, A. L., Erickson, K. E., Choudhury, A., Pines, G., and Gill, R. T. (2016) The Resistome: a comprehensive database of *Escherichia coli* resistance phenotypes. ACS Synthetic Biology 5, 1566–1577.

14. Erickson, K. E., Winkler, J. D., Nguyen, D. T., Gill, R. T., and Chatterjee, A. (2017) The Tolerome: A Database of Transcriptome-Level Contributions to Diverse *Escherichia coli* Resistance and Tolerance Phenotypes. ACS Synthetic Biology 6, 2302–2315.

15. Carbonell, P., Jervis, A. J., Robinson, C. J., Yan, C., Dunstan, M., Swainston, N., Vinaixa, M., Hollywood, K. A., Currin, A., and Rattray, N. J. (2018) An automated Design-Build-Test-Learn pipeline for enhanced microbial production of fine chemicals. Communications Biology 1, 66.

16. Camacho, D. M., Collins, K. M., Powers, R. K., Costello, J. C., and Collins, J. J. (2018) Next-Generation Machine Learning for Biological Networks. Cell

17. Marcellin, E., and Nielsen, L. K. (2018) Advances in analytical tools for high throughput strain engineering. Current Opinion in Biotechnology 54, 33–40.

18. Furusawa, C., Horinouchi, T., and Maeda, T. (2018) Toward prediction and control of antibiotic-resistance evolution. Current Opinion in Biotechnology 54, 45–49.

19. Caspi, R., Billington, R., Ferrer, L., Foerster, H., Fulcher, C. A., Keseler, I. M., Kothari, A., Krummenacker, M., Latendresse, M., Mueller, L. A., Ong, Q., Paley, S., Subhraveti, P., Weaver, D. S., and Karp, P. D. (2015) The MetaCyc database of metabolic pathways and enzymes and the BioCyc collection of pathway/genome databases. Nucleic Acids Research 44, D471–D480.

20. Krzywinski, M. I., Schein, J. E., Birol, I., Connors, J., Gascoyne, R., Horsman, D., Jones, S. J., and Marra, M. A. (2009) Circos: An information aesthetic for comparative genomics. Genome Research

21. Nichols, R. J. et al. (2011) Phenotypic Landscape of a Bacterial Cell. Cell 144, 143–156.

22. Zankari, E., Allesøe, R., Joensen, K. G., Cavaco, L. M., Lund, O., and Aarestrup, F. M. (2017) PointFinder: a novel web tool for WGS-based detection of antimicrobial resistance associated with chromosomal point mutations in bacterial pathogens. Journal of Antimicrobial Chemotherapy 72, 2764–2768.

23. Hecht, M., Bromberg, Y., and Rost, B. (2015) Better prediction of functional effects for sequence variants. BMC Genomics 16, S1.

24. Fariselli, P., Martelli, P. L., Savojardo, C., and Casadio, R. (2015) INPS: predicting the impact of non-synonymous variations on protein stability from sequence. Bioinformatics 31, 2816–2821.

25. Gama-Castro, S. et al. (2015) RegulonDB version 9.0: high-level integration of gene regulation, coexpression, motif clustering and beyond. Nucleic Acids Research 44, D133–D143.

26. Kim, H., Shim, J. E., Shin, J., and Lee, I. (2015) EcoliNet: a database of cofunctional gene network for Escherichia coli. Database 2015, bav001.

27. Fuhrer, T., Zampieri, M., Sévin, D. C., Sauer, U., and Zamboni, N. (2017) Genomewide landscape of gene–metabolome associations in *Escherichia coli*. Molecular Systems Biology 13, 907.

28. Deatherage, D. E., Kepner, J. L., Bennett, A. F., Lenski, R. E., and Barrick, J. E. (2017) Specificity of genome evolution in experimental populations of *Escherichia coli* evolved at different temperatures. PNAS 114, e1904–E1912.

29. Kusumawardhani, H., Hosseini, R., and de Winde, J. H. (2018) Solvent Tolerance in Bacteria: Fulfilling the Promise of the Biotech Era? Trends in Biotechnology

30. Chen, Y., and Nielsen, J. (2016) Biobased organic acids production by metabolically engineered microorganisms. Current Opinion in Biotechnology 37, 165–172.

31. Labrie, S. J., Samson, J. E., and Moineau, S. (2010) Bacteriophage resistance mechanisms. Nature Reviews Microbiology 8, 317.

32. Lang, G. I., Rice, D. P., Hickman, M. J., Sodergren, E., Weinstock, G. M., Botstein, D., and Desai, M. M. (2013) Pervasive genetic hitchhiking and clonal interference in forty evolving yeast populations. Nature 500, 571.

33. Reyes, L. H., Almario, M. P., Winkler, J., Orozco, M. M., and Kao, K. C. (2012) Visualizing evolution in real time to determine the molecular mechanisms of n-butanol tolerance in *Escherichia coli*. Metabolic Engineering 14, 579–590.

34. Jako?ciūnas, T., Pedersen, L. E., Lis, A. V., Jensen, M. K., and Keasling, J. D. (2018) CasPER, a method for directed evolution in genomic contexts using mutagenesis and CRISPR/Cas9. Metabolic Engineering 48, 288–296.

35. Halperin, S. O., Tou, C. J., Wong, E. B., Modavi, C., Schaffer, D. V., and Dueber, J. E. (2018) CRISPR-guided DNA polymerases enable diversification of all nucleotides in a tunable window. Nature 1.

36. Kuznetsov, G., Goodman, D. B., Filsinger, G. T., Landon, M., Rohland, N., Aach, J., Lajoie, M. J., and Church, G. M. (2017) Optimizing complex phenotypes through model-guided multiplex genome engineering. Genome Biology 18, 100.

37. Si, T., Chao, R., Min, Y., Wu, Y., Ren, W., and Zhao, H. (2017) Automated multiplex genome-scale engineering in yeast. Nature Communications 8, 15187.

38. Tenaillon, O., Rodrίguez-Verdugo, A., Gaut, R. L., McDonald, P., Bennett, A. F., Long, A. D., and Gaut, B. S. (2012) The molecular diversity of adaptive convergence. Science 335, 457–461.

39. Rodrίguez-Verdugo, A., Tenaillon, O., and Gaut, B. S. (2015) First-step mutations during adaptation restore the expression of hundreds of genes. Molecular Biology and Evolution 33, 25–39.

40. Hug, S. M., and Gaut, B. S. (2015) The phenotypic signature of adaptation to thermal stress in *Escherichia coli*. BMC Evolutionary Biology 15, 177.

41. González-González, A., Hug, S. M., Rodrίguez-Verdugo, A., Patel, J. S., and Gaut, B. S. (2017) Adaptive mutations in RNA polymerase and the transcriptional terminator Rho have similar effects on *Escherichia coli* gene expression. Molecular Biology and Evolution 34, 2839–2855.

42. Mans, R., Daran, J.-M. G., and Pronk, J. T. (2018) Under pressure: Evolutionary engineering of yeast strains for improved performance in fuels and chemicals production. Current Opinion in Biotechnology 50, 47–56.

43. Zeitoun, R. I., Garst, A. D., Degen, G. D., Pines, G., Mansell, T. J., Glebes, T. Y., Boyle, N. R., and Gill, R. T. (2015) Multiplexed tracking of combinatorial genomic mutations in engineered cell populations. Nature Biotechnology 33, 631.

44. Warsi, O. M., Andersson, D. I., and Dykhuizen, D. E. (2018) Different adaptive strategies in *E. coli* populations evolving under macronutrient limitation and metal ion limitation. BMC Evolutionary Biology 18, 72.

45. Warner, J. R., Reeder, P. J., Karimpour-Fard, A., Woodruff, L. B., and Gill, R. T. (2010) Rapid profiling of a microbial genome using mixtures of barcoded oligonucleotides. Nature Biotechnology 28, 856–862.

46. Chan, Y., Chan, Y. K., Goodman, D. B., Guo, X., Chavez, A., Lim, E. T., and Church, G. M. (2018) Enabling multiplexed testing of pooled donor cells through whole-genome sequencing. Genome Medicine 10, 31.

47. Lee, H., Popodi, E., Tang, H., and Foster, P. L. (2012) Rate and molecular spectrum of spontaneous mutations in the bacterium *Escherichia coli* as determined by whole-genome sequencing. PNAS 109, E2774–E2783.

48. Pines, G., Winkler, J. D., Pines, A., and Gill, R. T. (2017) Refactoring the genetic code for increased evolvability. MBio 8, e01654–17.

49. Pines, G., Bassalo, M. C., Oh, E. J., Choudhury, A., Garst, A. D., and Gill, R. T. (2018) Isolation of Genomic Deoxyxylulose Phosphate Reductoisomerase (DXR) Mutations Conferring Resistance to Fosmidomycin. bioRxiv 296954.

50. Firnberg, E., and Ostermeier, M. (2013) The genetic code constrains yet facilitates Darwinian evolution. Nucleic acids research 41, 7420–7428.

51. Shoichet, B. K., Baase, W. A., Kuroki, R., and Matthews, B. W. (1995) A relationship between protein stability and protein function. PNAS 92, 452–456.

52. Winkler, J. D., Garcia, C., Olson, M., Callaway, E., and Kao, K. C. (2014) Evolved osmotolerant *Escherichia coli* mutants frequently exhibit defective N-acetylglucosamine catabolism and point mutations in cell shape-regulating protein MreB. Applied and Environmental Microbiology 80, 3729–3740.

53. Dillon, M. M., Rouillard, N. P., Van Dam, B., Gallet, R., and Cooper, V. S. (2016) Diverse phenotypic and genetic responses to short-term selection in evolving *Escherichia coli* populations. Evolution 70, 586–599.

54. Ma, J., Yu, M. K., Fong, S., Ono, K., Sage, E., Demchak, B., Sharan, R., and Ideker, T. (2018) Using deep learning to model the hierarchical structure and function of a cell. Nature Methods 15, 290.

55. LeCun, Y., Bengio, Y., and Hinton, G. (2015) Deep Learning. Nature 521, 436.

56. Rutherford, B. J., Dahl, R. H., Price, R. E., Szmidt, H. L., Benke, P. I., Mukhopadhyay, A., and Keasling, J. D. (2010) Functional genomic study of exogenous n-butanol stress in *Escherichia coli*. Applied and Environmental Microbiology 76, 1935–1945.

57. Rodrίguez-Verdugo, A., Gaut, B. S., and Tenaillon, O. (2013) Evolution of *Escherichia coli* rifampicin resistance in an antibiotic-free environment during thermal stress. BMC Evolutionary Biology 13, 50.

58. Costanzo, M., VanderSluis, B., Koch, E. N., Baryshnikova, A., Pons, C., Tan, G., Wang, W., Usaj, M., Hanchard, J., Lee, S. D., and Pelechano, V. (2016) A global genetic interaction network maps a wiring diagram of cellular function. Science 353, aaf1420.

59. Deatherage, D. E., and Barrick, J. E. Engineering and analyzing multicellular systems; Springer, 2014; pp 165–188.

60. Hansen, A. S. L., Lennen, R. M., Sonnenschein, N., and Herrgård, M. J. (2017) Systems biology solutions for biochemical production challenges. Current Opinion in Biotechnology 45, 85–91.

